# Vocal error monitoring in the primate auditory cortex

**DOI:** 10.1101/2024.06.15.599151

**Authors:** Steven J Eliades, Joji Tsunada

**Affiliations:** Department of Head and Neck Surgery & Communication Sciences, Duke University School of Medicine, Durham, NC, USA; Chinese Institute for Brain Research, Beijing, China

## Abstract

Sensory-motor control requires the integration and monitoring of sensory feedback resulting from our behaviors. This self-monitoring is thought to result from comparisons between predictions of expected sensory consequences of action and the feedback actually received, resulting in activity that encodes feedback error. Although similar mechanisms have been proposed during speech and vocal production, including sensitivity to experimentally-perturbed auditory feedback, evidence for a vocal ‘error signal’ has been limited. Here, we recorded from the auditory cortex of vocalizing non-human primates, using real-time frequency shifts to introduce feedback errors of varying magnitude and direction. We found neural activity that scaled with the magnitude of feedback error in both directions, consistent with vocal error monitoring at both the individual unit and population levels. This feedback sensitivity was greater than that predicted based upon passive sensory responses and was more specific for units in the vocal frequency range. Similar patterns of sensitivity were seen in response to natural variations in produced vocal acoustics. These results provide evidence that the auditory cortex encodes the degree of vocal feedback error using both unit-level error calculations and changes in the population of neurons involved. These mechanisms may provide critical error information necessary for feedback-dependent vocal control.

## Introduction

A fundamental question in systems neuroscience is how does the brain deal with self-generated sensory inputs and, in particular, how does the brain encode differences between the predictable consequences of our actions and the sensory feedback it actually receives? It has been proposed that a common underlying mechanism is shared between different species and different sensorimotor systems in which a predictive signal, termed a corollary discharge, is relayed from motor areas to sensory ones ^1–10^. These signals are thought to contain predictions about the expected sensory responses during motor activity, which can then be directly compared to sensory feedback in order to determine errors in motor output, and are basis of models of feedback-dependent motor control ^11–14^. Despite the intuitive appeal of these models, direct evidence for such ‘error signals’ has been, surprisingly, limited.

Vocal production is an important sensory-motor behavior that is common amongst many animal species and shares many features typical of sensorimotor systems. Importantly, accurate vocal communication requires auditory self-monitoring to detect and correct production errors^15^ or errors in sound transmission due to the acoustic environment ^16,17^. Consistent with sensorimotor models, there is a well-described suppression of the auditory cortex during both human speech ^18–25^ and animal vocal production ^26–30^. When experimental perturbations introduce a mismatch between the expected vocal sounds produced and sensory feedback received, there is an increase in cortical activity that is enhanced compared to similar manipulations during passive listening ^30–32^. This increased sensory activity (or loss of suppression) predicts subsequent compensatory changes in vocal production, and may be the driver of feedback-dependent vocal error correction ^22,33–35^. It remains unclear, however, whether or not this feedback sensitivity is consistent with the classic notion of an ‘error signal,’ i.e. a neural response that scales with magnitude of sensory feedback error. With a few exceptions^36–38^, the majority of previous speech/vocal studies in both humans and animals have been limited to a single feedback manipulation and, as a result, it is unknown whether responses to altered feedback are encoding a specific sensory mismatch, or represent a non-specific response to the manipulation. It is also unclear how such a putative error calculation might be implemented, whether it is a population-level phenomenon or is evident at the level of individual neurons.

We therefore sought to determine whether individual neurons in the auditory cortex exhibited feedback sensitivity features consistent with a vocal error signal. We recorded cortical neurons in vocalizing marmoset monkeys (*Callithrix jacchus*) while altering auditory feedback using frequency shifts of varying magnitude and direction, comparing results to passive sensory responses. We found a sensitivity to altered vocal feedback at the population level that scaled with magnitude of the feedback error, with individual units exhibiting varying sensitivity to feedback in a frequency tuning-dependent manner. We also found that neural activity was sensitive to natural variability in vocal acoustics, though less than during altered feedback, suggesting that vocal sensory prediction may be relatively course. Together these results are consistent with vocal feedback error signal calculation occurring at both the unit and population level.

## Results

We recorded the activities of 4351 units from the auditory cortexes of three marmoset monkeys while the animals made voluntary, self-initiated vocalizations, comparing results between normal vocal production and vocalizations with frequency-shifted acoustic feedback. Figure 1A illustrates a sample unit during normal and altered vocal feedback. This unit was suppressed during normal vocalization, but exhibited increased firing during frequency shifted feedback, particularly during −2 semitone (ST) shifts where activity actually exceeded baseline. Consistent with previous work ^27,39,40^, we found the majority of auditory cortical units exhibited suppression during normal vocalizations, including the presence of decreased firing prior to the onset of vocal production (Fig. 1B,C). When animals vocalized in the presence of frequency-shifted feedback, most units responded with increases in their firing rates, more evident in those units that were most suppressed. At the population level, these feedback responses were similar for positive and negative frequency shifts (Fig. 1C). Comparing population responses for feedback shifts of varying magnitude and direction (−3 to 3 ST) revealed increasing average responses with the degree of feedback shift (error), evident for shifts in both directions (Fig. 1D,E). Overall, there was a significant population correlation with the magnitude of the feedback shift (r=0.16, p<0.001). This feedback sensitivity increased with the magnitude of normal, baseline vocal suppression, seen for feedback shifts of different magnitudes and directions (Fig. 1F, r=-0.26, p<0.001), consistent with previous work suggesting that vocal feedback responses were stronger in units exhibiting vocal suppression than those that were not suppressed^30,33,41^. We further compared the effects of feedback shift magnitude to the degree of normal vocal suppression, and found larger feedback effects for suppressed units (Fig. 1G). Collectively these results suggest that population activity exhibits feedback responses that depends on the magnitude of feedback error, present for both directions of error, consistent with the features of an error signal at the population level.

**Figure 1:**
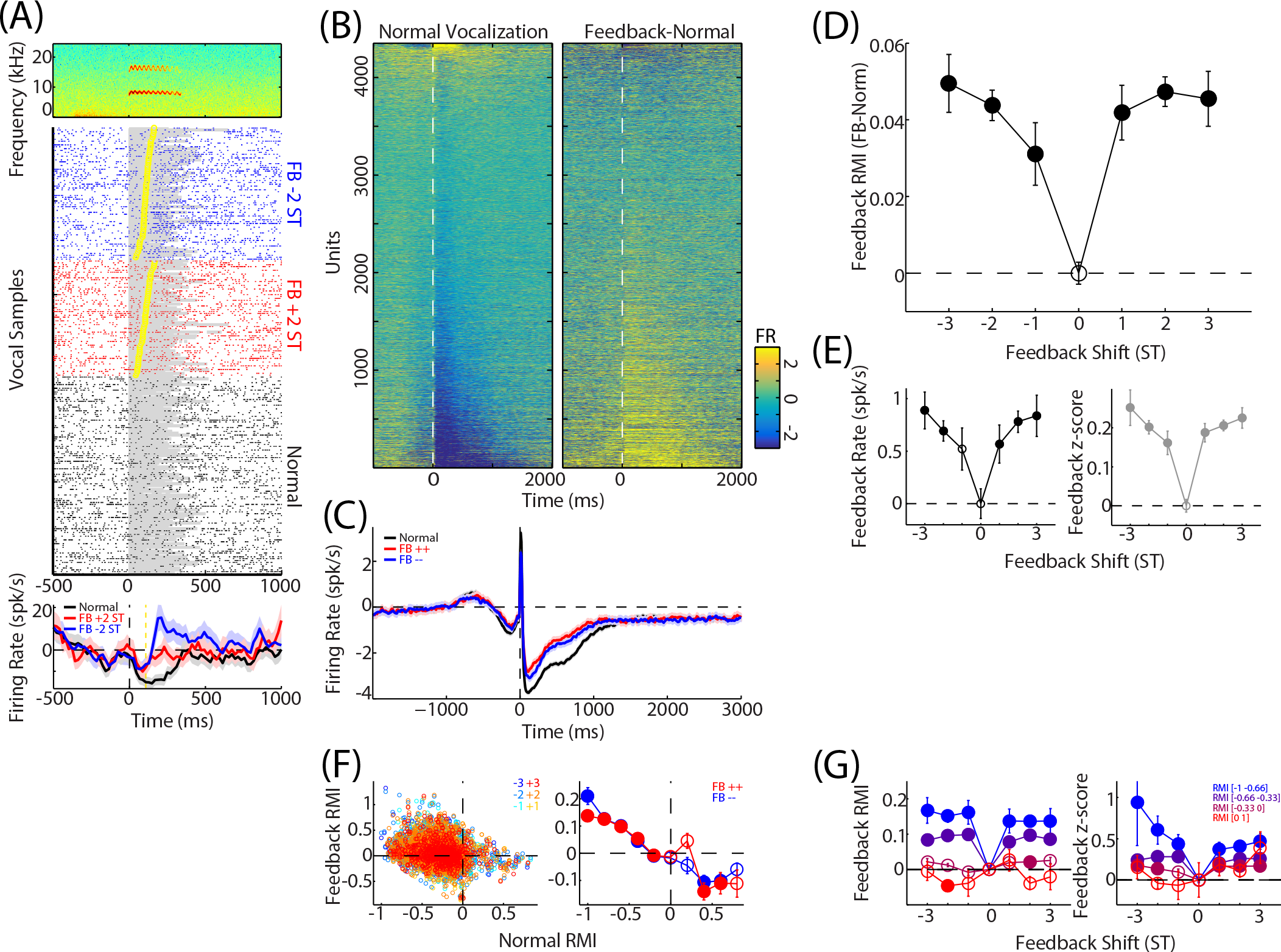
Auditory cortex units are sensitive to frequency-shifted vocal feedback. (A) A sample unit exhibiting suppression during normal vocalization and sensitivity to both positive and negative feedback (‘FB’) shifts. Spectrogram of a trill vocalization is shown (top), as well as raster plots (middle), and vocal onset-aligned PSTHs (bottom, mean and SEM) during normal and shifted vocalizations. Durations of individual vocalizations are indicated (shaded) as well as onset times for feedback shifts (yellow markers, average: yellow line). (B) Population heatmap plot of normal vocal responses from the unit population (left), ordered from most suppressed to most excited. Changes due to shifted feedback (right) for the same units exhibited increased activity, particularly for units with greater suppression. (C) Population average PSTHs illustrating vocalization-induced suppression during normal vocalization and average increased firing for both positive and negative feedback changes. Mean and 95% confidence intervals are shown. (D) Comparisons of population average feedback effects (feedback - normal Response Modulation Index [RMI]) between different feedback shifts revealed activity that increased with the magnitude of the perturbation, seen for shifts in both directions. Zero point illustrates the variability of normal vocal responses. Mean and SEM are shown; filled symbols: p<0.05. (E) Alternative normalization using raw firing rate changes (left) and z-scored firing rates are also shown (right). (F) Comparisons of feedback responses with normal vocal activity revealed stronger feedback effects in more suppressed units, shown for both individual shifts (left) and averaged into positive and negative groups (right). (G) Average effects of different feedbacks, divided by units’ normal vocal responses. Both z-score (left) and RMI (right) normalized feedback measures are shown. Stronger feedback responses, and greater scaling with feedback magnitude, were seen in more suppressed units.

In order to determine to what degree this error-like population feedback sensitivity could have been a simply a result of underlying sensory responses, instead of depending on a motor input, we examined how feedback responses compared to frequency tuning and passive vocal playback. Using both tone and band-pass noise stimuli, we measured unit center frequencies (CFs) and found that feedback sensitivity was highly dependent on CF, with the strongest effects of altered feedback for units with CFs in the vocal frequency range (6-16 kHz including both fundamental and the first harmonic), and weaker feedback responses in lower CF units (Fig. 2A,B). Interestingly, there was also a subset of units in much higher frequencies which also exhibited feedback responses, particularly for positive feedback shifts. Overall, however, we did not see a large difference between positive and negative shifts. We also noted CF dependence of baseline vocal suppression, with most units exhibiting a degree of vocal suppression and larger suppression for units in the vocal range (Fig. 2A). Similarly, the scaling of feedback responses with the magnitude of altered feedback was also stronger in the vocal frequency range (Fig. 2C,D).

**Figure 2:**
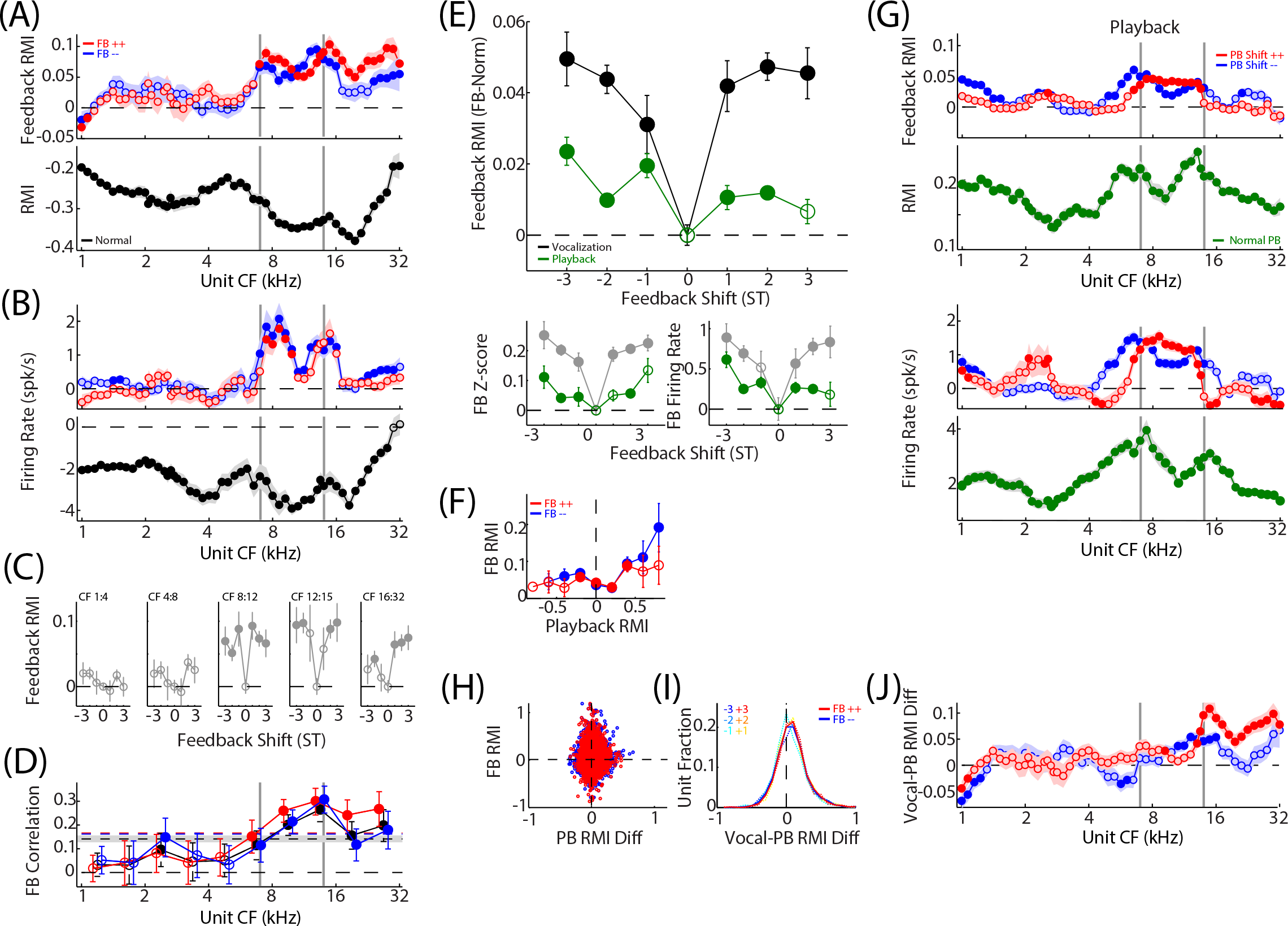
Feedback sensitivity is frequency-dependent and greater than passive sensory responses. (A) Average feedback RMI responses are shown, sorted based by unit center frequency (CF), grouped into positive and negative shifts. Stronger feedback effects were seen in the vocal frequency range (grey bars: 7 kHz vocal fundamental [f0] and harmonic) and for some units with CFs above the typical range. Both positive and negative shifts evoked feedback responses over this range, but no specific pattern for shift direction was evident, except a positive shift bias above the vocal range. Normal vocal responses (bottom) also exhibited stronger vocal suppression over this same vocal frequency range. Mean and SEM (shaded) are shown, filled symbols p<0.05. Similar patterns of responses were seen for feedback effects using un-normalized firing rates, though feedback responses were notably more CF constrained than for RMI (B). (C) Feedback RMI responses are shown, comparing different feedback magnitudes and grouped based upon unit CFs. Scaling of responses with feedback was more evident in higher CF units, though some vocal-range units showed sensitivity to feedback that did not further increase with the feedback magnitude. (D) Correlation coefficients measured between feedback RMI and feedback magnitude, and plotted for units of different CF, were larger for units in the vocal range. Correlations were separately measured for positive and negative feedback as well as the average of both directions; global population correlations are indicated (horizontal lines). Although this correlation increased in parallel with strength of feedback responses, there was a difference between correlations near vocal f0 and the harmonic, with stronger correlations near the later, even though feedback responses were similar for both. (E) Comparing feedback effects to responses evoked by similar frequency shifts during passive vocal playback showed larger effects during vocal production. Mean and SEM are shown (filled: p<0.05) and results are separately plotted for effects measured using RMI (top), z-score (bottom left), and firing rates (bottom right). (F) Vocal feedback effects are plotted against normal, un-shifted playback, showing larger feedback effects in units with stronger sensory responses to vocal sounds. (G) Shifted and normal playback responses are plotted against unit CF for both RMI (top) and firing rate (bottom) measures. Like vocal production, playback responses were CF dependent, including during normal playback, though a larger range of unit CFs were responsive to the feedback shift than seen during vocal production. Playback also showed differences between positive and negative frequency shifts that were not seen during production. (H) Scatter plot directly comparing frequency shift effects between vocal production and playback, showing larger feedback effects during vocal production for both RMI (left) and firing rate (right), but little correlation. (I) Histogram of RMI differences between vocal production and playback, suggesting an increase in feedback sensitivity (pos shifts: 0.026±0.22, p<0.001; neg shifts: 0.034±0.22, p<0.001). (J) Comparison of feedback production-playback differences for units of different CF. Differences were larger in higher frequencies.

Comparing vocal feedback responses with those evoked during passive listening to vocal sounds, we found larger frequency shift effects during vocal production, though passive responses also tended to scale with shifts (Fig. 2E). Feedback effects were larger in units with stronger playback responses (Fig. 2F), consistent with a model in which coding during vocal production integrates inputs from the ascending auditory pathway with suppressive top-down corollary discharge effects. Not unexpectedly, examining the CF dependence of playback responses also showed stronger responses to playback, with and without frequency shifts, for units in the vocal frequency range (Fig. 2G). Interestingly comparing frequency shifts during vocal production and playback (i.e. Fig. 2A-B vs. Fig. 2G) revealed that vocal activity was more CF constrained than for playback, particularly in low CF units, suggesting that one effect of corollary discharges may be to narrow the population-level tuning to those units more specifically sensitive to the vocal range. Directly comparing frequency shift responses during vocal production and playback within individual units revealed greater feedback sensitivity during vocal production (Fig. 2H-I), consistent with previous results^30^. These sensitivity increases were actually strongest for units with CFs outside the vocal frequency range, primarily on the higher side, and more modest for those within (Fig 2J). Collectively these results suggest that putative suppressive corollary discharge mechanisms during vocal production target auditory cortex units in a frequency-dependent fashion, while simultaneously selectively enhancing sensitivity to feedback error beyond what could be explained by passive sensory mechanisms.

While these results demonstrate population-level sensitivity to feedback error, it is unclear whether this finding is a result of feedback coding within the firing rates of individual units or whether population-level feedback responses could instead emerge through the recruitment of additional neurons. For example, one possible model could be a corollary discharge that selectively or differentially suppresses units that best respond to expected vocal acoustic feedback, with frequency shifted feedback instead activating adjacent CF units, resulting in a population-average increase. These two alternatives would be fundamentally different mechanisms of sensory-motor integration. We therefore examined error coding within individual units, using different combinations of feedback magnitude and direction. Figure 3A illustrates an example unit comparing responses to both positive and negative feedback shifts (±2 ST) and showing similar responses in both directions. Other units showed differential responses, with preferences for either positive or negative feedback errors (Fig. 3B,C). Averaging across all units tested with both ±2 ST shifts we found symmetric population responses (Fig. 3D). Direct comparisons between shifts showed strong correlations (r=0.89, p<0.001,), though there was variability between units (Fig. 3E). Overall, 43% of units showed significant responses to at least one feedback shift: 15% selectively responded to one shift or the other (7% negative, 8% positive), while 28% were significantly responsive to both directions (Fig. 3F). A larger fraction of units were sensitive to feedback in the right hemisphere compared to the left, and in more medial recording electrode (putatively primary auditory cortex; χ^2^=15.1, p=0.002 and χ^2^=20.7, p=0.014, respectively). Differences between ±2 shifts were widely distributed between units, with a non-significant bias (+2 vs. −2) towards negative shifts (Fig. 3G). There was no systematic variation in the direction of shift bias/preference with the degree of vocal suppression, however suppressed units were more likely to have (absolute value) bias for on direction over another (Fig. 3H), suggesting that more suppressed units were not only more responsive to feedback, but were also more likely to show differences between different feedback conditions. Comparing feedback bias direction between vocal production and playback revealed no systematic relationship (Fig. 3I), however units with any playback bias were also likely to have bias during vocal production, with a slight increase in the magnitude of the absolute bias during vocal production (Fig J-K). Examining bias direction and absolute magnitude for units of different CFs did not show a systematic relationship, though units in the vocal frequency range showed slightly larger average bias magnitude (Fig. 3L). More focused examination of shift direction bias around vocal fundamental (f0) and harmonic frequencies revealed preferences for negative shifts in units below vocal frequencies, and preferences for positive shifts in units just above vocal frequencies (Fig. 3M). Overall, we were more likely to find units with significant feedback responses in the vocal frequency range (Fig. 3N, χ^2^=77.8, p<0.001). These results suggest that many individual units were sensitive to feedback error, though this feedback sensitivity varied between different units, in part dependent on normal vocal suppression, playback responses, and frequency tuning.

**Figure 3:**
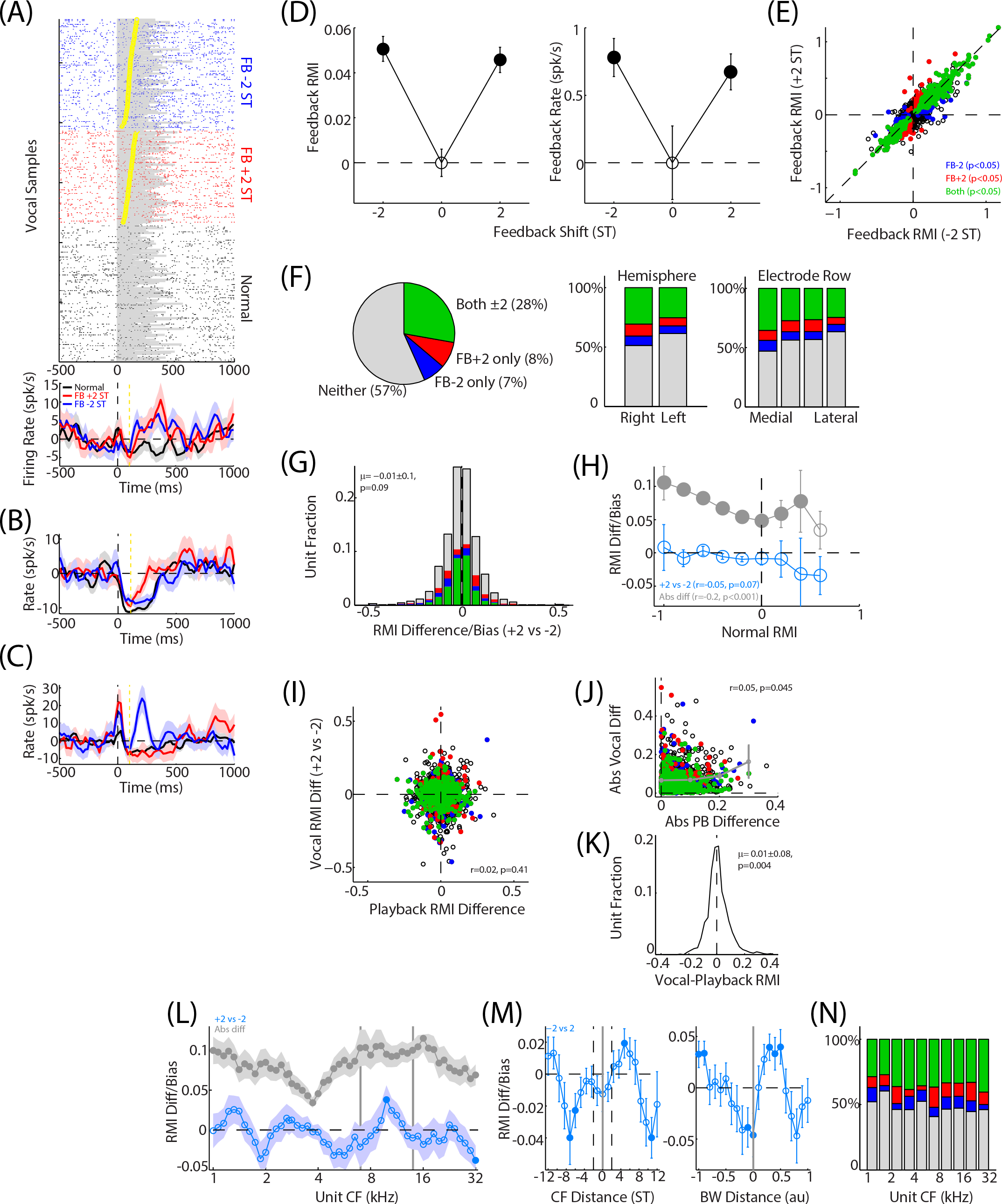
Comparisons of feedback direction effects within individual units. (A) A representative sample unit is shown, including raster (top) and PSTH (bottom), with similar responses to both +2 and −2 ST feedback shifts. (B,C) Additional sample units showing sensitivity to only one of the feedback directions. (D) Average feedback responses (mean±SEM) were similar for both feedback directions (±2 ST) for both RMI and firing rate measures. Filled p<0.05. (E) Raster plot directly comparing feedback direction effects. Units are colored by significant responses for either or both feedback directions. Distribution of units with significant feedback responses are shown (F), compared between right and left hemisphere (top) and different electrode array rows (medial to lateral, bottom). (G) Histogram comparing feedback direction bias for individual units (+2 response minus −2), showing no significant bias for one feedback direction over another. Significant units have been colored. Comparing direction bias against normal vocal suppression did not show any trends, suggesting overall balance for ±2 ST responses, however the absolute value of the direction bias was larger in more suppressed units (H). (I) Scatter plot comparing feedback direction bias between vocal production and playback, showing little correlation. Units with significant feedback effects (colored as in E) showed differences in both directions. Comparisons of absolute value bias showed larger vocal production effects in units with larger absolute bias during playback (J), and overall greater absolute bias during vocal production (K). (L) Plot of feedback direction bias and absolute bias against unit CF suggesting small differences for frequencies straddling vocal f0. Absolute bias was only slightly stronger in the vocal frequency range. (M) Plot of feedback direction bias against the octave distance between unit CF and reference vocal f0 (or harmonic), shown in semitone distance. Units with CFs below f0 had average bias towards −2 ST (negative direction bias), while units above f0 had positive biases. The 2 ST feedback range is indicated (vertical dashed lines). Similar results were seen when CF distance was normalized by unit bandwidth (right). Overall, more units with significant feedback responses were seen in the vocal frequency range (N).

We also tested a number of units with different combinations of frequency shift magnitudes (+1 vs. +2 ST, +2/+3, +2/+4, +1/+2/+3, −1/−2, −2/−3, −2/−4, and −1/−2/−3). Figure 4A shows a sample unit which was responsive to +4 ST feedback shifts, but not to +2 ST, while Fig 4B illustrates a unit in which both −2 and −4 ST shifts evoked activity, but more so for the larger shift. Examining averages responses to these various feedback combinations showed activity that, in general, increased with the magnitude of the feedback error (Fig. 4C-E). There was considerable variability in some shift combinations, possibly due to variability between the unit subsets tested, and in particular the effects of normalization, notably evident in units tested in −1/−2/−3 ST which showed feedback responses by firing rate and z-score, but not when RMI normalized. Quantifying the effects of feedback error magnitude revealed larger correlations between feedback response and shift magnitude (i.e. 0/+2/+3), as well as differences between individual shifts (+2 vs. +3), particularly for larger shift magnitudes (Fig. 4F). These feedback error measures (shift magnitude correlations and differences between individual shift pairs) correlated with the degree of normal vocal suppression for many of the shift pairs tested, though shift difference comparisons were not significant (Fig. 4G). Comparing these responses to similar frequency shifts during passive playback, we again found greater sensitivity during vocal production (Fig. 4H-I). Shift magnitude correlations were larger for units with stronger average playback responses, but there was little relationship between production and playback for individual shift differences (Fig. 4I). As with population averages, feedback error correlations for individual units were dependent on CF, with largest correlations in the vocal frequency range (Fig. 4J). Interestingly, the CF range of shift correlations increased with the magnitude of the feedback shift, suggesting recruitment of additional units to encode feedback, and correlations often reversed outside the vocal range. We did not find any systematic relationship between individual shift differences and unit CF (Fig. 4K). We also noted that the same shifts often evoked quite different responses depending upon the context in which they were delivered (such as +2 in the +2/+4 and +2/+3 testing, Fig 4C-E). Whether these differences were a result of different unit sets, or some larger contextual modulation remains an intriguing question. Together with the ±2 ST results, these findings suggest that individual units are, in fact, sensitive to feedback magnitude and direction, consistent with feedback error encoding at both the individual and population level, but this encoding is specific for subsets of neurons that are tuned to the vocal frequency range and respond to vocal sounds during passive playback.

**Figure 4:**
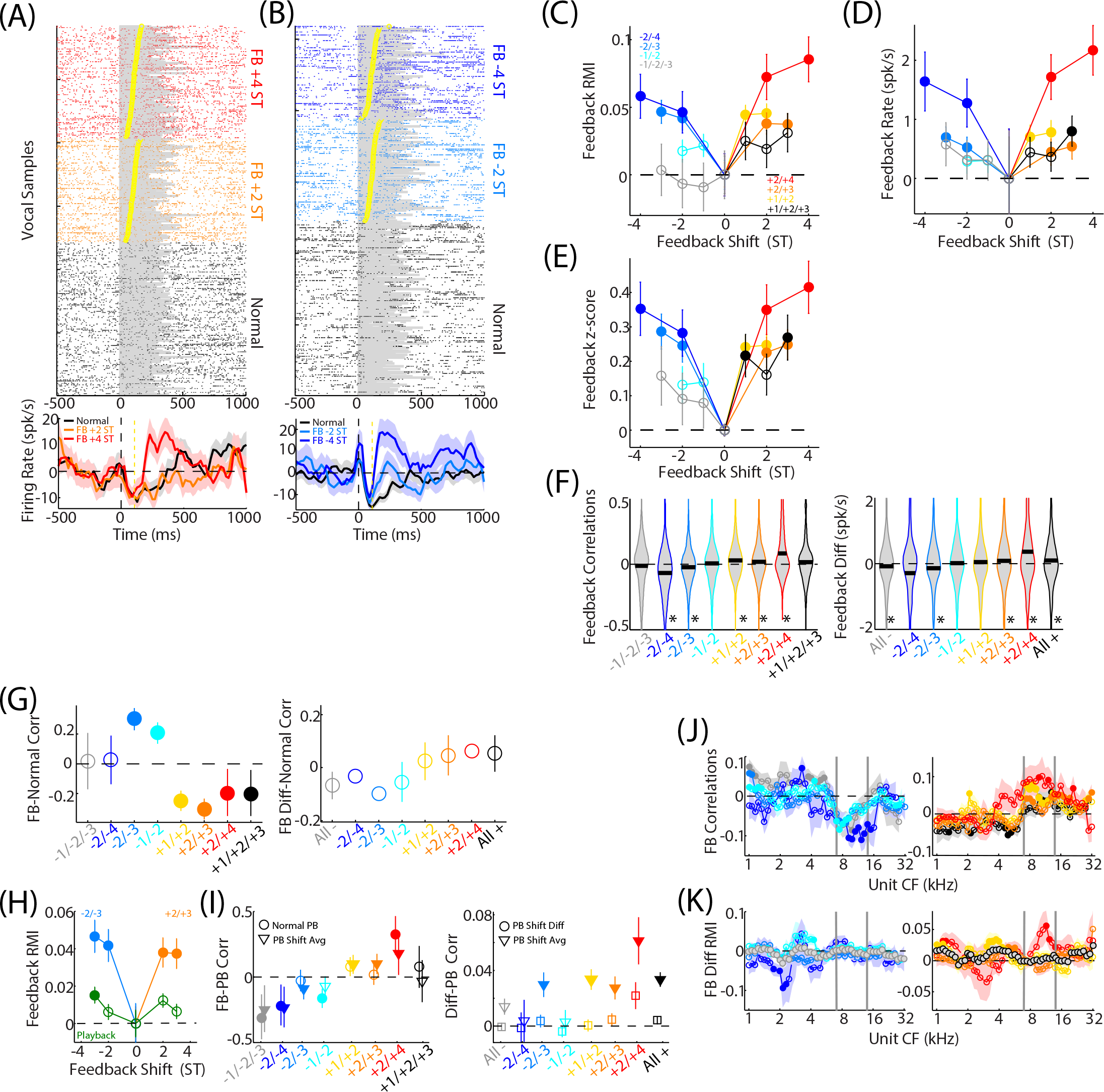
Individual units show sensitivity for increasing feedback magnitude. (A) An example unit is shown, demonstrating increased responses to +4 but not +2 ST feedback shifts. Another sample unit, studied with −2 and −4 ST feedback showed responses to both, but exhibited greater firing rates for the larger shift (B). (C-E) Comparisons of feedback magnitude and vocal responses for a variety of tested shift sets (mean±SEM, filled p<0.05). Results are shown separately when measured using RMI (C), firing rates (D), and z-score (E) normalization. In general responses were larger with increasing feedback magnitude, though a few feedback comparisons did not exhibit any changes or feedback sensitivities. Interestingly, ±2 ST feedback shifts, the most commonly tested, exhibited different responses depending on the feedback set. (F) Feedback effects within individual units were quantified using a correlation coefficient between firing rate and feedback magnitude (including zero/no feedback), and direct RMI comparison between the different feedbacks (i.e. +2 vs +3 ST). Positive correlations/differences indicate increased activity with a positive shift comparison, while negative indicated increased activity with a negative shift comparison (i.e. a −3 vs −2 ST differences). Histograms are shown illustrating the distribution of these measurements, with significant correlations and differences noted primarily for larger shifts (* p<0.05). Averaged feedback differences are also included, grouping all positive and negative shifts (right). (G) Feedback magnitude correlations (left) and differences for individual shift pairs (right) were compared with the suppression seen during normal vocalization and correlated across the unit population. A significant population correlation was noted for most feedback magnitude comparisons, suggesting stronger feedback effects for more suppressed units (left, filled: p<0.05). Feedback pair comparisons showed trends suggesting larger differences in more suppressed units, but results were not significant (p>0.05). (H) Comparison of +2/+3 and −2/−3 ST frequency shifts between vocal production and playback, showing larger effects during vocal production. (I) Unit vocal feedback correlations and pair differences were compared to playback and plotted as a population correlation coefficient with both normal playback and average playback shift responses (left). Stronger feedback effects, for both positive and negative FB directions, were seen for units with greater playback responses. Vocal production FB differences were similarly stronger in units with larger playback shift responses, but did not significantly correlate with differences between the different playback shifts (right). (J) Comparing feedback correlations with unit CFs showed significant results primarily in the vocal frequency range, and often inverted correlations for units lower than vocal frequencies. Results are shown separately for negative (left) and positive (right) shift sets. (K) Feedback pair differences plotted against unit CF did not show any systematic relationships.

These findings of frequency-specific feedback error encoding give rise to another important question as to the nature and specificity of the putative predictive corollary discharge. Are predictions specific to expected acoustics for a single phoneme or vocalization, or are they a more general prediction for averaged acoustical output? Directly measuring corollary discharge signals has thus far been challenging, however recent work in human speech has found MEG activity that varied with vowel formant variability, particularly with larger deviations from the mean, suggested to represent an error response compared from an average rather than individual template^23^. Past work in marmosets has also suggested some variability in vocal suppression with vocal acoustics^40^. We therefore examined how responses during normal, un-altered, vocal production changed with vocal acoustic variability and compared results to responses during frequency-shifted feedback. Figure 5A illustrates a sample unit that was strongly suppressed during vocalization, but where the degree of suppression was correlated with the mean fundamental frequency (f0) of the vocalizations. Measuring correlations coefficients between neural activity and f0 across the unit population revealed a wide distribution, though with a net bias towards positive correlations, increasing firing rate with f0 (Fig. 5B). Because such acoustic-response relationships could also potentially vary in an error-like fashion, i.e. with distance from the mean and not just absolute f0, we also measured correlations separately for vocalizations above and below the mean f0 (measured for individual animals and call types). Vocalizations with acoustics below the mean (‘lower half’) had negative average correlations between activity and f0, suggesting increasing activity with decreasing f0, while vocalizations above the mean (‘upper half’) had net positive correlations (Fig. 5B). This increased activity as acoustics move away from the mean suggest a response pattern similar to observed ‘V-shaped’ error responses during frequency-shifted feedback, though there was a bias towards positive correlations. Overall, 13.8% of units exhibit significant raw acoustic correlations (n=189), while 8.5% and 12.8% units had significant lower and upper half correlations (n=102/169). Similar analysis for vocal loudness (SPL) did not result in a similar correlation pattern. There was also no systematic relationship between upper and lower half f0 correlations within individual units (Fig. 5C). We examined raw vocal acoustic correlations and compared results to acoustic correlations measured during passive playback for units of different CFs (Fig. 5D). Unsurprisingly, vocal acoustic correlations were strongest for units in the vocal frequency range, particularly focused around vocal mean f0. Interestingly however, the set of units with acoustic correlations was broader during playback and more focused during vocal production, even though both had matched f0 ranges. Comparing upper and lower half correlations showed similar CF-dependence, with stronger correlations in both directions for units matching vocal frequencies, and more variable effects outside this range (Fig. 5E). Because correlations normalize the magnitude of responses, we also examined linear regression coefficients and compared vocal production and playback to determine if the strength of responses changed along with the CF tuning (Fig. 5F,G). Although more units had significant acoustic regression/correlation during playback compared to vocal production (25.4% vs 13.8%), the magnitude of the regression coefficients were much larger during vocal production, but did not correlate (r=0.03, p=0.35). These results show that the dependence of unit responses on vocal acoustics was stronger during vocal production than playback, and was present in a more CF-specific group of units. These results also suggest that there is an error-like response even with normal vocal acoustic variability that cannot be explained based upon sensory responses alone, and may suggest that vocal predictions are not specific for individual vocalizations produced.

**Figure 5:**
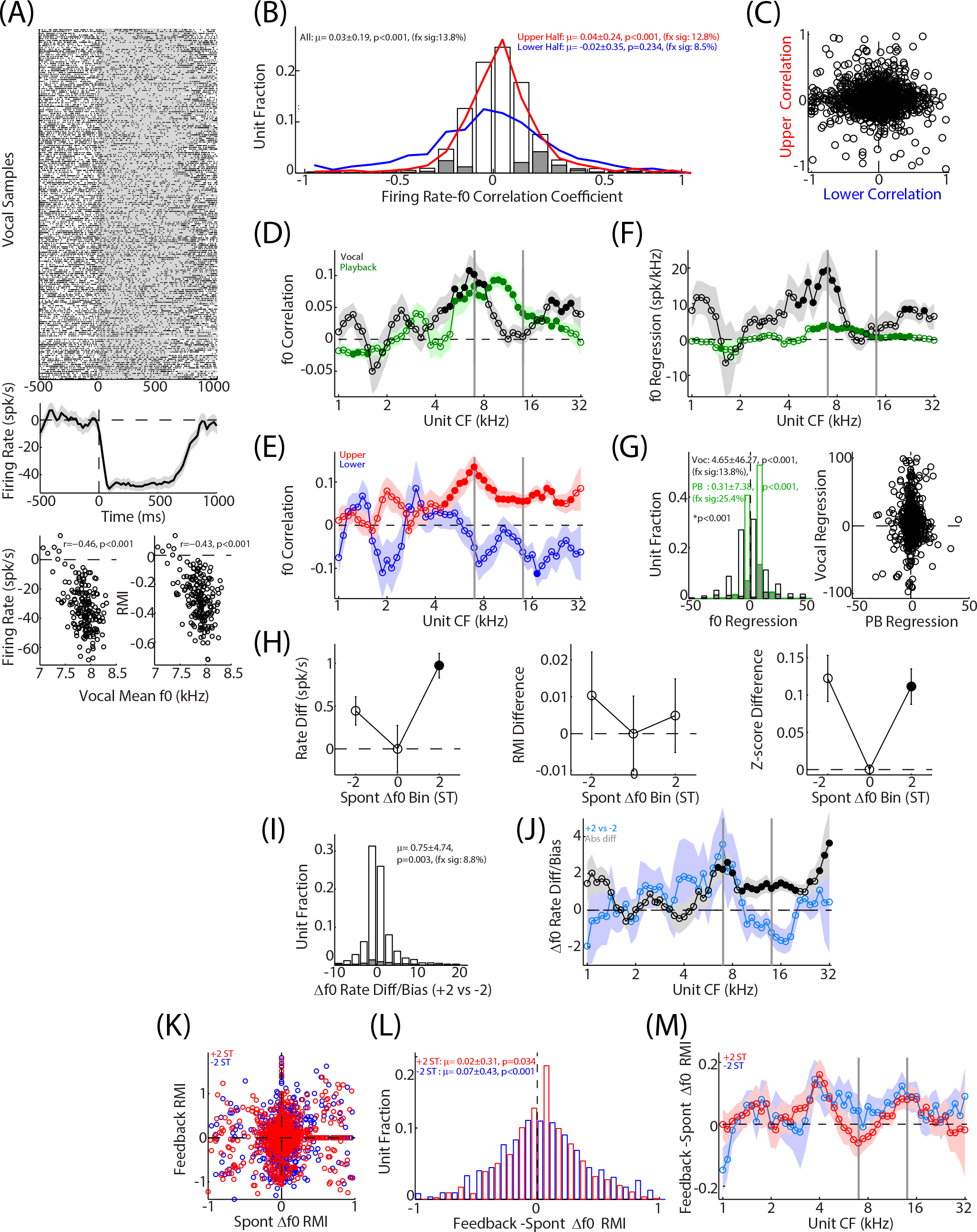
Vocal responses change with natural vocal variability. (A) Sample unit showing suppressed responses during normal (un-shifted) vocal production, including raster (top) and PSTH (middle). Individual trials correlated with variations in vocal f0, measured with both firing rate (bottom left) and RMI (bottom right). (B) Histogram showing distribution of correlation coefficients between firing rate and vocal f0 across the unit population, illustrating a net bias towards positive correlations. Filled bars indicate units with significant correlations (p<0.05). Correlations were also separately calculated for vocalizations above and below the mean vocal f0 and are overlayed on the distribution plot. Results measured on the lower half had a bias towards negative correlations (increased firing with decreasing f0), while the upper half showed positive frequency correlations. (C) Comparison of upper and lower half correlations within individual units, showing little relationship. (D) Comparison of f0 correlations between units of different CFs, and between vocal production and playback. Both showed a bias towards positive average correlation, however these were present for units across the whole vocal frequency range during playback, but only around the mean vocal f0 during vocal production. (E) CF comparison of upper and lower half correlations, showing symmetric results around vocal frequencies. (F) Linear regression coefficients between firing rate and vocal f0, compared by CF and between vocal production and playback. Coefficients were larger during vocal production, particularly around vocal frequencies. (G) Direct comparisons of production and playback regression as histograms (left) and scatter plot (right), showing significantly larger coefficients during vocal production. (H) Mean vocal production responses measured for natural vocal variability with f0 matching ±2 ST feedback shifts, revealed patterns similar to those seen during feedback experiments, shown for firing rates (left), RMI (middle) and z-scored (right) measures. (I) Histogram comparing responses in +2 and −2 matched f0 bins. Responses were larger in +2, consistent with the measured bias towards positive f0 correlations. (J) Comparing mean responses in the ±2 ST bins, and their difference, across CFs showed larger responses in the vocal frequency range. (K) Comparison of RMI differences during shifted feedback trials with effects of natural f0 variation showed weak correlations for positive but not negative shifts (+2/upper: r=0.13, p<0.001; −2/lower: r=0.05, p=0.08), however histograms of the difference (L) revealed larger changes during experimental feedback shifts in both directions. (M) Differences between responses to experimental shifts and natural variability are plotted against unit CF. Enhancement for the experimental feedback perturbation was larger for units outside the traditional vocal frequency range, both above and below, and more similar to natural variations in the middle of the vocal range.

In order to compare the effects of natural vocal acoustic variability with frequency-shifted feedback, we measured responses to normal vocalizations divided into categories based upon individual vocalization f0: vocal mean (spanning −0.5 to +0.5 ST range compared to the mean), −2 ST (span: −2.5 to −1.5 ST), and +2 ST (+1.5 to 2.5 ST). Responses measured in these categories showed increased activity in ±2 ST bins compared to vocal mean, with a slight bias towards +2 (Fig. 5 H-I), consistent with the correlation results (i.e. Fig. 5B). Examining the CF dependence for mean ±2 ST responses, and their differences, showed largest effects in the vocal f0 range, as noted with other measures (Fig. 5J). Directly comparing shifted feedback and natural variability revealed that experimentally-perturbed feedback resulted in larger responses than for vocal variability, even when the acoustic changes from the mean/normal f0 were the same (Fig. 5K-L). Differences were larger for units CFs outside the typical vocal f0 range, with increases most notable for units around 4 and 16 kHz, and closer matches around the 7-8 kHz vocal f0 range. These results further support the suggestion that spontaneous vocal variability around an average template evokes an error-like response, but that the effects of experimentally-induced errors are greater, particularly for more frequency distant units. It should be noted, however, that our analysis of shifted feedback responses did not take into account variability that might exist within the acoustics of shifted vocal trials, and could lead to variability within the feedback results.

## Discussion

In this study, we examined feedback error coding in the auditory cortex of vocalizing non-human primates. Using frequency-shifted feedback of varying magnitude and direction, we found responses that increased with the magnitude of the feedback mismatch in both positive and negative shift directions. This feedback sensitivity was present at both the population and individual unit level, was greater than predicted from frequency shifts during passive listening, and was dependent on unit frequency tuning. We also found that natural variations in vocal acoustics evoked similar changes to experimentally altered feedback, though to a lesser degree. Collectively, these auditory cortical responses exhibit characteristics suggestive of a vocal ‘error signal’, a response when there is a mismatch between expected sensory input and the actual feedback received, notably a suppression of responses when error is minimal, i.e. no feedback changes, and an increase in neural responses as the degree of mismatch increases.

Sensitivity to altered feedback has been previously demonstrated during both human speech and animal vocalization ^22,25,30,32–34^. However, most studies have been limited to a single feedback manipulation. As a result, it has been unclear if feedback sensitivity was truly encoding feedback error, or was just a non-specific increase in sound-evoked activity. A small number of human speech studies have compared parametrically-varied feedback and found increased responses with the magnitude of the feedback ^36–38^. Our results show similar findings, but also exhibited symmetric sensitivities for feedback changes in both positive and negative directions, an important feature to establish that neural responses are encoding feedback error and not simply vocal acoustics. However, given that it is also important to be able to perceive the direction of feedback error, and to make the appropriate compensatory behavioral responses^33^, how the brain is able to make this distinction is unclear. Although population-level feedback responses were symmetric in both directions, and many individual units were also sensitive to both feedback directions, a small number of units did exhibit selective direction preferences, and these selective units may be a potential mechanism by which feedback direction, and resulting behavioral response, can be encoded.

It remains unclear as to the specificity of sensory predictions used in calculating any putative vocal errors. Based upon models of feedback error coding and feedback-dependent vocal control, one might expect a corollary discharge signal relaying very specific predictions about expected vocal acoustics for comparison with sensory feedback ^12,42^. However, recent studies using natural speech formant variability found that there were differences between neural responses to speech tokens that were more central or average vs. speech acoustics that were further from the mean ^23^. We found similar results, with correlations between neural activity and the deviation in vocal production from an animal’s mean acoustics. These results suggest that sensory prediction might be somewhat coarse, relaying predictions about average vocal acoustics, rather than specific events or vocal features, or that predictions are based upon an early, less developed motor plan and that specific acoustics are generated later in brain areas that introduce a degree of noise or variability, as seen in other motor systems^43^. This sort of average prediction may have certain advantages, notably dealing with time varying acoustics during the duration of vocal production, where attempting to encode error on a moment-to-moment basis, combined with delays in sensory processing, might lead to instability in feedback-dependent vocal control ^44–47^. A coarser prediction may also be potentially useful in the setting of noisy, variable vocal control. There may also be an advantage in simplifying the neural connectivity needed to relay specific vocal acoustics predictions from motor to sensory areas.

We observed increased sensitivity to vocal feedback mismatch when compared to similar frequency shifts during passive listening. Previous work has also found increased feedback sensitivity during vocal production in both marmosets and humans ^30–32^. Although we noted overall stronger responses to shifted feedback during vocal production, on average vocal responses did not do a better job distinguishing different feedback manipulations than would be expecte based on passive coding. There are several possible explanations for this apparent contradiction. The degree of frequent shift (1-4 semitones) is small relative to the size of auditory cortex receptive fields, and therefore analysis based on individual units may be insensitive to these differences, and discrimination may only emerge at a population level. We did note that there was some increased sensitivity in differentiating different feedback, but primarily for neurons on the edge of the vocal frequency range. It is possible that one effect of the corollary discharge is to enhance feedback error coding in these border neurons that might otherwise be relatively insensitive to the frequency shift. Finally, it also possible that the trial-to-trial variability in vocal acoustics, which can sometimes exceed the feedback shifts, masks the benefits of sensory prediction.

Another novel observation in these studies is the frequency-dependence of vocal feedback responses. While we have previously noted that normal vocal suppression can vary between units based upon CF^29,41^, with greater suppression in the vocal range, we had not previously found variation in feedback sensitivity with unit CF. Here we noted that altered feedback responses were stronger in units in the vocal frequency range. This is, perhaps, not surprising. Putatively, corollary discharge predictions are being compared to sensory feedback through the ascending auditory pathway. We would not expect neurons well outside the range of vocal acoustics to receive much in the way of auditory input during vocal production. Similar ranges of sensitivity were noted for passive playback of vocal sounds, both with and without frequency shifts. Interestingly, however, was that the range of CF dependence was different between vocal production and playback, both in population and unit level feedback responses as well as in the effects of spontaneous vocal variability, with vocal production exhibiting more focused feedback responses. While this could have been a result of slightly different vocal acoustics between playback and production, the matched CF peaks during unmodified vocalization/playback suggests a relatively good acoustic matches. It is possible that the effects of corollary discharges not only selectively suppress some sets of units, but also selectively enhance feedback responses for units in the vocal feedback range.

Looking more broadly, how do sensory-motor systems use corollary discharge predictions to encode sensory feedback errors. One could envision two possible extremes to these models. Under a rate-based model, different neurons receive specific predictions about vocal acoustics, and individually encode feedback mismatch. This sort of model is consistent with most broader notions about predictive coding in sensory systems ^48–50^. An alternative model, based upon population coding, could encode feedback error by selectively inhibiting only those neurons expected to respond to normal vocalizations and then activating additional neurons in adjacent CF regions with increasing frequency-shift feedback. Both models are compatible with existing human speech data. The present experiments show elements of both forms of vocal feedback coding. Individual units exhibited responses that scaled with the magnitude of feedback error, both experimentally induced and from spontaneous vocal variability. These results suggest that these units receive individual corollary discharge predictions. At the same time, there is a degree of CF specificity for the suppression during normal vocal responses, and greater feedback sensitivity in nearby CF units, suggesting that these corollary discharge signals may also be selective in the neurons they innervate. However, given the limited range of vocal feedback acoustics and manipulations, it is still possible that units outside the vocal frequency range might be responsive to feedback error when subjected to appropriate vocal acoustics.

Results of these studies have potentially important implications for understanding feedback error coding and vocal control. The ability to encode vocal feedback errors is important for feedback-dependent vocal control^33,51^, which may be an evolutionarily ancient precursor to sensorimotor learning of human speech. Corollary discharge predictions have also been linked to our ability to discriminate between self-generated and external sounds^52^, as well as to detect unexpected acoustics form non-vocal self-generated sounds ^53–56^. Dysfunction in this error coding has also been implicated in vocal communication disorders, including stuttering ^57–61^. Finally, dysfunctional corollary discharge predictions, and resulting aberrant self-monitoring, has been implicated in the auditory hallucinations in schizophrenia ^62–64^. More broadly, these results demonstrate how feedback error can be encoded at both the neuronal and population level, which has implications for multiple other prediction-based sensory-motor systems.

## Methods

We recorded neural activities from three adult marmoset monkeys (*Callithrix jacchus*), one female and two males, while the animals produced self-initiated vocalizations. Neural activity from the auditory cortex, including both primary and non-primary areas, was recorded using implanted multi-electrode arrays and compared to simultaneously recorded vocal behavior. A subset of units with +2 and −2 semitone (ST) responses have been included in a previous publication^33^, however no direct comparisons of the feedback were done in those studies. All experiments were conducted under the guidelines and protocols approved by the University of Pennsylvania Care and Use Committee, where the studies were conducted.

### Vocal Recordings and Altered Feedback

Using previous methods, we recorded marmosets vocalizing while in their home colony^30,33,65^. Animals were placed in a small cage within a custom three-walled sound attenuating booth allowing free visual and vocal interaction with the rest of the marmosets in the colony. During recordings, marmosets were tethered within a small cage, to allow neural recording, but were otherwise unrestrained. Vocalizations were recorded using a directional microphone (AKGC1000S) placed ∼20cm in front of the animal and digitized at 48.8 kHz sampling rate (TDT RX-8, Tucker-Davis Technologies, Alachua FL). Additional colony-facing microphones were used to discriminate vocalizations from the test animal from the rest of the colony. Vocalizations were extracted from recordings and spectrographically classified into established marmoset call types^66^ using a semi-automated system. All major call types were produced in this context, however only long-duration calls (phee, trillphee, trill) were included in this analysis. Acoustic analyses were performed on the vocalizations, using the spectrogram to measure the vocal fundamental frequency and then averaging across the call duration, as well as measurements of peak and average sound-pressure level (SPL). Other acoustic features, including harmonics or specific spectrotemporal features were not measured.

During altered feedback experiments, microphone signals were passed through a digital effects processor (Eventide Eclipse V4), shifted either up or down in frequency, and presented back to the animal at +10dB through modified ear-bud style headphones (Sony MDR-EX10LP). Altered feedback was presented during a random subset of vocalizations (p ∼ 0.5-0.8) with an average 100ms delay from vocal onset under a control of a computer that detected vocal onset and randomly selected the feedback shift to deliver. In order to get sufficient trials for the units being recorded, on any given day only a subset of feedback shifts were tested (either +2/−2, +1/+2, +2/+3, +2/+4, −1/−2, −2/3, −2/−4, +1/+2/+3, or −1/−2/−3 ST). Due to limited number of vocalizations and sessions, some feedback combinations (notable +1/+2/+3, −1/−2/−3, +2/+4, and −2/−4) were tested less regularly. While animals wore headphones throughout the duration of a vocal recording sessions, these did not occlude the ear canal and were not connected to any sound sources during a period of baseline ‘normal’ vocal production at the start of each session.

### Neural Recordings

All marmosets were implanted bilaterally with multi-electrode arrays (Warp 16, Neuralynx, Bozeman MT), one in each auditory cortex. Details of the array design and recording technique have been previously published^65^. These arrays consist of a 4×4 grid of individually moveable sharp microelectrodes (4 MΩ tungsten; FHC, Bowdoinham ME). As in our previous methods, we first localized the center of primary auditory cortex using single-electrode methods, and placed arrays to cover the full range of the tonotopic axis, verified by frequency tuning. Based upon relative responses to tone and noise stimuli, electrodes were judged to span both primary (A1) and non-primary (belt, parabelt) auditory cortex^67^. Neural signals were observed on-line to guide electrode and optimize signal quality. Digitized signals were sorted off-line using custom MATLAB software and a principle component-based clustering method, and then classified as either single-unit or multi-units^65^. We included both single- and multi-unit responses in our analyses.

### Auditory Stimuli

Prior to each neural recording in the colony, we first presented auditory stimuli to characterize sensory tuning properties of the auditory cortex units. Marmosets were seated in a custom primate chair within a soundproof chamber (Industrial Acoustics, Bronx NY). Auditory stimuli were digitally generated at 97.6 kHz sampling rate and delivered using TDT hardware (System III) in free-field through a speaker (B& 686 S2) located ∼1 m in front of the animal. Tuning stimuli included tones (1-32 kHz, 10/octave; −10 to 80 dB SPL by 10 dB) and bandpass noise (1-32 kHz, 5/octave, 1 octave bandwidth), as well as wide-band noise stimuli. The center frequency (CF) of a unit’s tuning was determined by the strongest response to either tone or bandpass stimuli. Vocal playback stimuli were samples of an animal’s own vocalizations (previously recorded from that animal) presented at multiple sound levels. Although vocal stimuli were presented at different sound levels, only those samples overlapping vocal production loudness were used for comparisons between vocal production and auditory playback. We performed frequency shifts on vocal samples to match the feedback delivered during vocal experiments using a phase vocoder approach^30,33^. When comparing to frequency-shifted vocal production experiments, where animals hear both shifted and unshifted versions of their vocalizations, shifted playback was created by adding together both shifted and unshifted samples with the same relative feedback onset delay (100ms) and amplitude (+10 dB). For comparisons to spontaneous vocal variation, shifted feedback samples were tested covering the same range of acoustics observed during the vocal production, but delivered without adding to normal samples.

### Data Analysis

All analyses were performed using MALAB. We calculated spontaneous-subtracted firing rates individually for each vocalization, and averaged across responses. For display purposes, vocal onset-aligned peri-stimulus time histograms (PSTHs) were calculated with 10ms binwidth and smoothed with a 5-point moving average. Confidence intervals were calculated based upon the response standard error (SEM) over time. Consistent with previous work, we further quantified different units’ responses using a normalized rate metric, the Response Modulation Index (RMI), RMI = (FR_voc_-FR_pre_)/(FR_voc_+FR_pre_), where FR_voc_ and FR_pre_ are the firing rates before and during vocalization^27^. Pre-vocal firing rates were calculated from a window from 4 seconds to 1 second preceding vocal onset in order to exclude the effects of pre-vocal suppression which has been seen in the 200-500 msec before vocal onset^27,39^. We corrected for changes in mean pre-vocal firing rate during frequency-shifted trials ^31,33^. Based upon previous work showing similar responses between different vocal calltypes^39^, we pooled all three major call types in these analyses. As a control, we repeated some analyses limited to trill vocalizations and found qualitatively similar responses. In some analyses, units were categorized as suppressed or excited based upon a vocal RMI (comparing phrases to pre-vocal baseline) of ≤-0.2 or ≥0.1, consistent with our previous work. Similar calculations were performed for vocal playback.

Frequency tuning analyses were performed by calculating the center frequency (CF) of a unit as the frequency with the maximal firing rate response. The strongest response from either tone or bandpass noise tuning was used. Bandwidth was defined as the frequency range over which responses were at least 50% of the peak firing rate. Population CF distributions were calculated by binning units by their CF, smoothed by grouping units with a 5-bin moving average, and then averaging responses from those units, For some analyses, a unit CF distance was calculated as the ratio of the unit CF relative to a reference frequency in the middle of the vocal f0 range (7.5 kHz), in octaves. For units with CF closer to the vocal harmonic than f0, the octave distance from the harmonic reference frequency was used instead. A normalized CF distance metric was also calculated by dividing the CF distance by the bandwidth.

For analysis based upon vocal acoustic variability, we first measured the mean frequency and loudness (SPL) for each vocal sample. Vocal acoustics were then normalized by dividing each vocalization’s measurements by the mean acoustics observed for that specific animal and vocal call type. We repeated this normalization using the mean acoustics measured for vocalizations from a given recording session, rather than averaged across all sessions for an animal, and found similar results. These normalized measures were used to determine the mean vocal acoustics for correlation analyses and to match the ±2 semitone ranges tested using altered feedback (ranges: [−2.5 to 1.5 ST] and [1.5 to 2.5 ST]).

With the exception of correlation and regression analyses, all statistical tests were performed using non-parametric methods. Wilcoxon rank-sum and signed-rank tests (two-sided) were used to test the differences between unmatched and matched distribution medians, respectively. Kruskal-Wallis ANOVAs were used when comparing more than two conditions. Significant unit responses were determined using rank-sum comparisons of RMI responses between normal vocalizations and shifted feedback. Correlation values within individual units, and between units, were calculated with Pearson correlation coefficients. When binning and averaging results in a single plot, p-values were first calculated for individual bins/phrases and then False Discovery Rate (FDR) corrected for multiple comparisons.

## References

1 Blakemore, S. J., Goodbody, S. J. & Wolpert, D. M. Predicting the consequences of our own actions: the role of sensorimotor context estimation. J Neurosci 18, 7511–7518. (1998).

2 Sperry, R. W. Neural basis of the spontaneous optokinetic responses produced by visual inversion. J Comp Physiol Psych 43, 482–489 (1950).

3 von Holst, E. & Mittelstaedt, H. Das Reafferenzprinzip: Wechselwirkungen zwischen Zentralnervensystem und Peripherie. Naturwissenschaften 37, 464–476 (1950).

4 Wolpert, D. M., Ghahramani, Z. & Jordan, M. I. An internal model for sensorimotor integration. Science 269, 1880–1882 (1995).

5 Wolpert, D. M. & Flanagan, J. R. Motor prediction. Curr Biol 11, R729–732 (2001).

6 Crapse, T. B. & Sommer, M. A. Corollary discharge circuits in the primate brain. Curr Opin Neurobiol 18, 552–557 (2008). 10.1016/j.conb.2008.09.017

7 Crapse, T. B. & Sommer, M. A. Corollary discharge across the animal kingdom. Nat Rev Neurosci 9, 587–600 (2008). 10.1038/nrn2457

8 Poulet, J. F. & Hedwig, B. A corollary discharge maintains auditory sensitivity during sound production. Nature 418, 872–876. (2002).

9 Ross, J., Morrone, M. C., Goldberg, M. E. & Burr, D. C. Changes in visual perception at the time of saccades. Trends Neurosci 24, 113–121 (2001). 10.1016/s0166-2236(00)01685-4

10 Sommer, M. A. & Wurtz, R. H. A pathway in primate brain for internal monitoring of movements. Science 296, 1480–1482. (2002).

11 Houde, J. F. & Chang, E. F. The cortical computations underlying feedback control in vocal production. Curr Opin Neurobiol 33, 174–181 (2015). 10.1016/j.conb.2015.04.006

12 Houde, J. F. & Nagarajan, S. S. Speech production as state feedback control. Frontiers in human neuroscience 5, 82 (2011). 10.3389/fnhum.2011.00082

13 Wolpert, D. M. & Miall, R. C. Forward models for physiological motor control. Neural Netw 9, 1265–1279 (1996).

14 Wolpert, D. M. & Kawato, M. Multiple paired forward and inverse models for motor control. Neural Netw 11, 1317–1329 (1998).

15 Levelt, W. J. Monitoring and self-repair in speech. Cognition 14, 41–104 (1983).

16 Luo, J., Hage, S. R. & Moss, C. F. The Lombard Effect: From acoustics to neural mechanisms. Trends Neurosci 41, 938–949 (2018). 10.1016/j.tins.2018.07.011

17 Hage, S. R., Jiang, T., Berquist, S. W., Feng, J. & Metzner, W. Ambient noise induces independent shifts in call frequency and amplitude within the Lombard effect in echolocating bats. Proc Natl Acad Sci U S A 110, 4063–4068 (2013). 10.1073/pnas.1211533110

18 Paus, T., Perry, D. W., Zatorre, R. J., Worsley, K. J. & Evans, A. C. Modulation of cerebral blood flow in the human auditory cortex during speech: role of motor-to-sensory discharges. Eur J Neurosci 8, 2236–2246 (1996).

19 Binder, J. R., Liebenthal, E., Possing, E. T., Medler, D. A. & Ward, B. D. Neural correlates of sensory and decision processes in auditory object identification. Nat Neurosci 7, 295–301 (2004).

20 Crone, N. E. et al. Electrocorticographic gamma activity during word production in spoken and sign language. Neurology 57, 2045–2053. (2001).

21 Greenlee, J. D. et al. Human auditory cortical activation during self-vocalization. PLoS One 6, e14744 (2011). 10.1371/journal.pone.0014744

22 Chang, E. F., Niziolek, C. A., Knight, R. T., Nagarajan, S. S. & Houde, J. F. Human cortical sensorimotor network underlying feedback control of vocal pitch. Proc Natl Acad Sci U S A 110, 2653–2658 (2013). 10.1073/pnas.1216827110

23 Niziolek, C. A., Nagarajan, S. S. & Houde, J. F. What does motor efference copy represent? Evidence from speech production. J Neurosci 33, 16110–16116 (2013). 10.1523/JNEUROSCI.2137-13.2013

24 Flinker, A. et al. Single-trial speech suppression of auditory cortex activity in humans. J Neurosci 30, 16643–16650 (2010). 10.1523/JNEUROSCI.1809-10.2010

25 Houde, J. F., Nagarajan, S. S., Sekihara, K. & Merzenich, M. M. Modulation of the auditory cortex during speech: an MEG study. J. Cog. Neurosci. 14, 1125–1138 (2002).

26 Muller-Preuss, P. & Ploog, D. Inhibition of auditory cortical neurons during phonation. Brain Res 215, 61–76 (1981).

27 Eliades, S. J. & Wang, X. Sensory-motor interaction in the primate auditory cortex during self-initiated vocalizations. J Neurophysiol 89, 2194–2207 (2003).

28 Eliades, S. J. & Wang, X. Corollary Discharge Mechanisms During Vocal Production in Marmoset Monkeys. Biological psychiatry. Cognitive neuroscience and neuroimaging 4, 805–812 (2019). 10.1016/j.bpsc.2019.06.008

29 Tsunada, J. & Eliades, S. J. Dissociation of Unit Activity and Gamma Oscillations during Vocalization in Primate Auditory Cortex. J Neurosci 40, 4158–4171 (2020). 10.1523/JNEUROSCI.2749-19.2020

30 Eliades, S. J. & Wang, X. Neural substrates of vocalization feedback monitoring in primate auditory cortex. Nature 453, 1102–1106 (2008).

31 Eliades, S. J. & Wang, X. Neural correlates of the lombard effect in primate auditory cortex. J Neurosci 32, 10737–10748 (2012). 10.1523/JNEUROSCI.3448-11.2012

32 Greenlee, J. D. et al. Sensory-motor interactions for vocal pitch monitoring in non-primary human auditory cortex. PLoS One 8, e60783 (2013). 10.1371/journal.pone.0060783

33 Eliades, S. J. & Tsunada, J. Auditory cortical activity drives feedback-dependent vocal control in marmosets. Nature communications 9, 2540 (2018). 10.1038/s41467-018-04961-8

34 Behroozmand, R. et al. Neural correlates of vocal production and motor control in human Heschl’s Gyrus. J Neurosci 36, 2302–2315 (2016). 10.1523/JNEUROSCI.3305-14.2016

35 Burnett, T. A., Freedland, M. B., Larson, C. R. & Hain, T. C. Voice F0 responses to manipulations in pitch feedback. J Acoust Soc Am 103, 3153–3161. (1998).

36 Behroozmand, R. & Larson, C. R. Error-dependent modulation of speech-induced auditory suppression for pitch-shifted voice feedback. BMC Neurosci 12, 54 (2011). 1471-2202-12-54 [pii] 10.1186/1471-2202-12-54

37 Behroozmand, R., Karvelis, L., Liu, H. & Larson, C. R. Vocalization-induced enhancement of the auditory cortex responsiveness during voice F0 feedback perturbation. Clin Neurophysiol 120, 1303–1312 (2009). S1388-2457(09)00340-X [pii] 10.1016/j.clinph.2009.04.022

38 Ozker, M., Doyle, W., Devinsky, O. & Flinker, A. A cortical network processes auditory error signals during human speech production to maintain fluency. PLoS Biol 20, e3001493 (2022). 10.1371/journal.pbio.3001493

39 Eliades, S. J. & Wang, X. Comparison of auditory-vocal interactions across multiple types of vocalizations in marmoset auditory cortex. J Neurophysiol 109, 1638–1657 (2013). 10.1152/jn.00698.2012

40 Eliades, S. J. & Wang, X. Dynamics of auditory-vocal interaction in monkey auditory cortex. Cereb Cortex 15, 1510–1523 (2005).

41 Tsunada, J., Wang, X. & Eliades, S. J. Multiple processes of vocal sensory-motor interaction in primate auditory cortex. Nature communications 15, 3093 (2024). 10.1038/s41467-024-47510-2

42 Hickok, G., Houde, J. & Rong, F. Sensorimotor integration in speech processing: computational basis and neural organization. Neuron 69, 407–422 (2011). 10.1016/j.neuron.2011.01.019

43 Churchland, M. M., Afshar, A. & Shenoy, K. V. A central source of movement variability. Neuron 52, 1085–1096 (2006). 10.1016/j.neuron.2006.10.034

44 Perkell, J. et al. Speech motor control: Acoustic goals, saturation effects, auditory feedback and internal models. Speech Com. 22, 227–250 (1997).

45 Perkell, J. S. et al. Time course of speech changes in response to unanticipated short-term changes in hearing state. J Acoust Soc Am 121, 2296–2311 (2007). 10.1121/1.2642349

46 Cai, S. et al. Weak responses to auditory feedback perturbation during articulation in persons who stutter: evidence for abnormal auditory-motor transformation. PLoS One 7, e41830 (2012). 10.1371/journal.pone.0041830

47 Houde, J. F. & Jordan, M. I. Sensorimotor adaptation in speech production. Science 279, 1213–1216. (1998).

48 Spratling, M. W. A review of predictive coding algorithms. Brain and cognition 112, 92–97 (2017). 10.1016/j.bandc.2015.11.003

49 Alink, A., Schwiedrzik, C. M., Kohler, A., Singer, W. & Muckli, L. Stimulus predictability reduces responses in primary visual cortex. J Neurosci 30, 2960–2966 (2010). 10.1523/JNEUROSCI.3730-10.2010

50 Walsh, K. S., McGovern, D. P., Clark, A. & O’Connell, R. G. Evaluating the neurophysiological evidence for predictive processing as a model of perception. Ann N Y Acad Sci 1464, 242–268 (2020). 10.1111/nyas.14321

51 Eliades, S. J. & Tsunada, J. Effects of Cortical Stimulation on Feedback-Dependent Vocal Control in Non-Human Primates. Laryngoscope 133 Suppl 2, S1–S10 (2023). 10.1002/lary.30175

52 Ford, J. M., Palzes, V. A., Roach, B. J. & Mathalon, D. H. Did I Do That? Abnormal Predictive Processes in Schizophrenia When Button Pressing to Deliver a Tone. Schizophrenia bulletin (2013). 10.1093/schbul/sbt072

53 Schneider, D. M. & Mooney, R. Motor-related signals in the auditory system for listening and learning. Curr Opin Neurobiol 33, 78–84 (2015). 10.1016/j.conb.2015.03.004

54 Schneider, D. M. & Mooney, R. How Movement Modulates Hearing. Annu Rev Neurosci 41, 553–572 (2018). 10.1146/annurev-neuro-072116-031215

55 Schneider, D. M., Sundararajan, J. & Mooney, R. A cortical filter that learns to suppress the acoustic consequences of movement. Nature 561, 391–395 (2018). 10.1038/s41586-018-0520-5

56 Audette, N. J., Zhou, W., La Chioma, A. & Schneider, D. M. Precise movement-based predictions in the mouse auditory cortex. Curr Biol 32, 4925–4940 e4926 (2022). 10.1016/j.cub.2022.09.064

57 Fairbanks, G. & Guttman, N. Effects of delayed auditory feedback upon articulation. J Speech Hear Disord 1, 12–22 (1958).

58 Gruber, L. Sensory feedback and stuttering. J Speech Hear Disord 30, 378–380 (1965).

59 Lee, B. S. Effects of delayed speech feedback. J Acoust Soc Am 22, 824–826 (1950).

60 Timmons, B. A. & Boudreau, J. P. Auditory feedback as a major factor in stuttering. J Speech Hear Disord 37, 476–484 (1972).

61 Postma, A. & Kolk, H. Error monitoring in people who stutter: evidence against auditory feedback defect theories. J Speech Hear Res 35, 1024–1032 (1992).

62 Ford, J. M. et al. Neurophysiological studies of auditory verbal hallucinations. Schizophrenia bulletin 38, 715–723 (2012). 10.1093/schbul/sbs009

63 Ford, J. M. & Mathalon, D. H. Electrophysiological evidence of corollary discharge dysfunction in schizophrenia during talking and thinking. J Psychiatr Res 38, 37–46 (2004).

64 Ford, J. M. et al. Neurophysiological evidence of corollary discharge dysfunction in schizophrenia. Am J Psychiatry 158, 2069–2071 (2001).

65 Eliades, S. J. & Wang, X. Chronic multi-electrode neural recording in free-roaming monkeys. J Neurosci Methods 172, 201–214 (2008).

66 Agamaite, J. A., Chang, C. J., Osmanski, M. S. & Wang, X. A quantitative acoustic analysis of the vocal repertoire of the common marmoset (Callithrix jacchus). J Acoust Soc Am 138, 2906 (2015). 10.1121/1.4934268

67 Rauschecker, J. P. & Tian, B. Processing of band-passed noise in the lateral auditory belt cortex of the rhesus monkey. J Neurophysiol 91, 2578–2589 (2004).

